# HIF-1α and HIF-2α transcription factors differentially regulate lung alveolar macrophage function

**DOI:** 10.1101/2025.11.25.690373

**Authors:** Elena Priego, Irene Adán-Barrientos, Ruth Conde-Garrosa, Sarai Martínez-Cano, Iria Sánchez, Diego Mañanes, Annalaura Mastrangelo, Joaquín Amores, Helena M. Izquierdo, David Sancho

## Abstract

Alveolar Macrophages (AMs) reside in the alveoli, where oxygen pressure is high, therefore maintaining an active degradation of hypoxia-inducible transcription factors (HIF) mediated by the Von Hippel-Lindau protein (pVHL) ubiquitin ligase complex. We previously found that *Vhl*-deficient AMs not sensing high oxygen pressure are immature and functionally impaired. Here we investigated the specific roles of HIF-1α and HIF-2α isoforms in the regulation of AM functions. With this aim, we combined deletion of *Vhl* with single or double deletion of *Hif1a* and/or *Hif2a* in AMs under the control of the CD11c promoter using a Cre-Lox system. Our work demonstrates that in *Vhl*-deficient macrophages, both HIF-1α and HIF-2α contribute to AM defective self-renewal, while HIF-2α plays a central role in regulating the impaired AM maturation associated with pVHL loss. HIF-1α promotes a glycolytic shift in alveolar macrophages, while HIF-2α hinders lipid oxidation and surfactant clearance. Thus, HIF-2α raises as a selective critical factor restraining the therapeutic potential of AMs to degrade surfactant excess in mice that have developed pulmonary alveolar proteinosis (PAP). Overall, regulation of both HIF-1α and HIF-2α isoforms is required for an optimal AM function, highlighting HIF-2α as a potential pharmacological target for secondary PAP.

## INTRODUCTION

Alveolar macrophages (AMs) are lung-resident cells maintained by self-renewal during homeostasis.^1^ Through the sensing of granulocyte-macrophage colony-stimulating factor (GM-CSF) ^2,3^ and transforming growth factor β (TGF-β),^4^ fetal liver monocytes differentiate into mature AMs in the lung alveoli during the first week of life. Mature AMs are essential for the regulation of surfactant in the lung, which is comprised of 90% lipids and is required to decrease surface tension. AMs have a remarkable ability for lipid catabolism and cholesterol handling,^5^ and these functions are sustained by enzymes and receptors whose expression is regulated by Liver-X family of receptors (LXRs), Peroxisome proliferator-activated family of receptors (PPARs), CCAAT enhancer-binding proteins (C/ EBPs), and sterol regulatory element-binding proteins (SREBPs) transcription factors.^6^ Changes in AM lipid metabolism are found in human respiratory diseases such as chronic obstructive pulmonary disease (COPD) ^6^ and pulmonary alveolar proteinosis (PAP).^7,8^ Mechanistically in mice, deletion of *Pparg*, a master regulator of lipid metabolism induced by GM-CSF, selectively impairs the differentiation of Ams.^3^ Depletion of the transcription factor TFAM, involved in mitochondrial biogenesis, leads to a reduction in AM numbers and impairs their maturation *in vivo.* ^5,9^ Overall, these studies have identified essential factors required for AM maturation and function and demonstrate that alterations of AM metabolism are associated to AM dysfunction. We have previously shown that pVHL is necessary for the adaptation of AMs to high oxygen levels upon birth, and that *Vhl* deletion renders immature and functionally impaired AMs, which also present an altered metabolic profile, including enhanced glycolysis and decreased lipid oxidation capacity.^10^ However, the role of each HIFα isoform in regulating the functional maturation of AMs has not been investigated.

pVHL is the substrate recognition unit of an E3 ubiquitin ligase complex involved in the ubiquitination and degradation of the alpha isoforms of hypoxia inducible factors (HIFs) in an oxygen dependent manner.^11–13^ HIFs are heterodimeric transcription factors that induce the expression of a set of genes functionally involved in the cellular adaptation to low oxygen tension or hypoxia, including the metabolic switch to glycolysis.^14,15^ HIFs are composed of an oxygen sensitive α subunit and a constitutive β subunit. Three α subunits have been described (HIF-1α, HIF-2α, and HIF-3α), which heterodimerize with HIF-1β subunit (also named Aryl Hydrocarbon Receptor Nuclear Translocator 1; ARNT1), forming HIF-1, HIF-2 and HIF-3 transcription factors, respectively.^16^ When dimerized with the HIF-1β subunit, HIF-1α and HIF-2α regulate the transcription of both shared and distinct target genes. HIF-3α acts as a negative regulator of HIF-1α and HIF-2α by competing for HIF-1β protein available for dimerization, although the HIF-3 dimer also induces the expression of specific target genes.^17^ Beyond HIFα regulation, pVHL plays a role in other cellular processes including cytokine signaling, regulation of senescence, formation of extracellular matrix and mitochondrial function.^18–25^ Notably, in our previous work, we found that genes related to the adaptation to hypoxia, including many targets of HIF-1α and HIF-2α, were downregulated within AM maturation after birth.^10^ Therefore, we hypothesize that HIF-1α and HIF-2α must be degraded to ensure the correct functional maturation of AMs in steady-state.

Here, we investigated the specific function of HIF-1α and HIF-2α in AM phenotypic maturation, self-renewal and surfactant clearance in the lung. To assess this, we generated four different strains of mice: CD11c-Cre *Vhl*^fl/fl^ (CD11cΔ*Vhl*), CD11c-Cre *Vhl^f^*^l/fl^*Hif1a*^fl/fl^ (CD11cΔ*VhlΔHif1a*), CD11c-Cre *Vhl^fl/fl^Hif2a^fl/^*^fl^ (CD11cΔ*VhlΔHif2a*) and CD11c-Cre *Vhl^fl/fl^Hif1a^fl/fl^Hif2a^fl/fl^*(CD11cΔ*VhlΔHif1aΔHif2a)*. We demonstrate that expression of either HIF-1α and HIF-2α alone is sufficient to intrinsically and differentially impair AM terminal maturation and self-renewal, with full rescue observed only in CD11cΔ*VhlΔHif1aΔHif2a* mice. Whereas HIF-1α drives AM metabolic shift towards glycolysis, HIF-2α disrupts lipid oxidation and surfactant clearance, and is the main isoform responsible for impaired AM maturation. Altogether, this study shows that HIF inactivation ensures the optimal maturation of AMs in the lung, regulating the metabolic adaptation to high oxygen concentration and highlights different functional specificities of each HIFα isoform.

## RESULTS

### HIF-1α is the driver of the metabolic switch to glycolysis in pVHL-deficient alveolar macrophages

Our previous work established that pVHL is required for AM phenotypic and functional maturation,^10^ but did not address the specific contribution of HIF-1α and HIF-2α or additional effects of pVHL. To analyze the specific function of HIF-1α and HIF-2α in AMs, we generated four different mouse strains: CD11c-Cre *Vhl*^fl/fl^ (CD11cΔ*Vhl*), CD11c-Cre *Vhl^f^*^l/fl^*Hif1a*^fl/fl^ (CD11cΔ*VhlΔHif1a*), CD11c-Cre *Vhl^fl/fl^Hif2a^fl/^*^fl^ (CD11cΔ*VhlΔHif2a*) and CD11c-Cre *Vhl^fl/fl^ Hif1a^fl/f^ ^l^Hif2a^fl/fl^* (CD11cΔ*VhlΔHif1aΔHif2a)*. Analysis of AMs from these mice revealed the expected genotype deficiencies in *Vhl*, *Hif1a*, and *Hif2a* expression (Figure S1A-C), with additional half reduction in *Hif1a* expression levels in CD11cΔ*VhlΔHif2a* mice (Figure S1B).

To investigate the contribution of HIF-1α and HIF-2α to the immature-like phenotype found in *Vhl*-deficient Ams,^10,26^ we performed an RNA-seq of AMs isolated from all the mice strains compared with their respective Cre negative littermate control mice. Analysis of differentially expressed genes (DEG) showed 1246 genes expressed with at least a 2-fold difference (Adj. p < 0.01) between each conditional KO and its control, except for CD11cΔ*Vhl*Δ*Hif1a*Δ*Hif2a* AMs, in which only *Vhl* and *Plcb4* were differentially expressed (Figure 1A-D), showing that the transcriptional changes seen in *Vhl*-deficient AM relies exclusively on downstream transcription factors HIF-1α and HIF-2α rather than on HIF-independent pVHL functions. Principal Component Analysis (PCA) showed that samples from CD11cΔ*Vhl*Δ*Hif1a*Δ*Hif2a* scored close to controls and distant from CD11cΔ*Vhl*, CD11cΔ*Vhl*Δ*Hif1a*, and CD11cΔ*Vhl*Δ*Hif2a* in principal component 1 (PC1) (Figure 1E).

**Figure 1.**
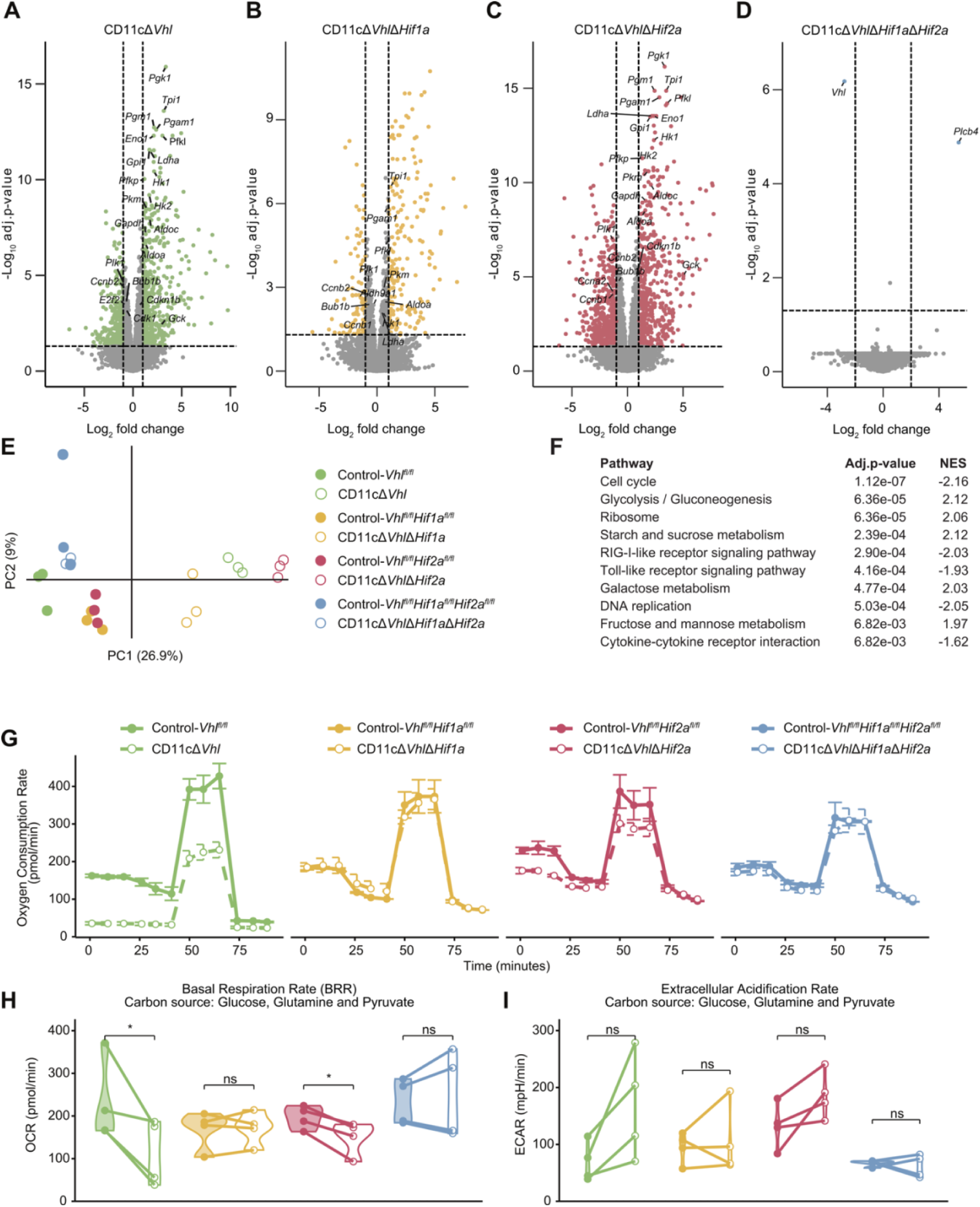
HIF1α drives glycolysis in VHL-deficient alveolar macrophages. (A-D) Volcano plot of differentially expressed genes ranked showing adjusted p-value (Log10 adj. p-value) and log2 fold change, between AMs from CD11cΔ*Vhl* (A), CD11cΔ*VhlΔHif1a* (B), CD11cΔ*VhlΔHif2a* (C) and CD11cΔ*VhlΔHif1aΔHif2a* (D) mice compared to their littermate control; differentially expressed genes (adjusted p-value < 0.05 and abs(log2FC) > 1) are depicted in the color code associated with each genotype. (E) Principal component analysis (PCA) of transcriptomic profiles of AMs from CD11cΔ*Vhl*, CD11cΔ*VhlΔHif1a*, CD11cΔ*VhlΔHif2a*, CD11cΔ*VhlΔHif1aΔHif2a* mice, and their littermate controls. Each dot represents a pull of 2-3 mice. (F) Pre-ranked enrichment analysis of the genes defining PC1 using KEGG Pathways. NES, normalized enrichment score; p-value adjusted with Benjamin–Hochberg correction. (G) Oxygen consumption rate (OCR) by BAL AMs cultured in media containing glucose, glutamine, and pyruvate. Data are shown as mean ± SD. (H, I) Basal respiratory rate (BRR) (H) and extracellular acidification rate (ECAR) (I) of BAL AMs. Each dot represents a pool of n = 6-7 mice per genotype. Paired Student’s t-test, *p<0.05 and “ns” stands for non statistically significant. See also Figures S1, S2.

Different genes were deregulated in AMs depending on the genotype (Figure S2A-C). Pre-ranked enrichment analysis of the genes with the greatest contribution to PC1 unveiled pathways related with cell division and metabolism, especially glucose related pathways, suggesting that both HIF-1α and HIF-2α are involved in their regulation (Figure 1F). To further interrogate the specific contribution of each HIF isoform, we performed an overrepresentation analysis, using the Gene Ontology (GO) database, of DEG from each genotype. Most of the downregulated pathways in *Vhl*-deficient AMs were restored upon further deletion of *Hif1a* and not *Hif2a*, such as cell cycle, DNA replication, p53 signaling pathway or fatty acid (FA) metabolism (Figure S2D). We also found pathways similarly controlled by HIF-1α and HIF-2α such as *AMPK signaling pathway* or *PI3K-Akt signaling pathway* (Figure S2D), suggesting that binding of either HIF isoform to the same gene promoters is sufficient to activate these pathways. In contrast, *PPAR signaling pathway*, *HIF1Α signaling pathway* or *Glycolysis/Gluconeogenesis* pathways were only partially restored by HIF-1α or HIF-2α deficiency, indicating their differential regulation by both transcription factors (Figure S2D).

HIF-1α and HIF-2α induce the transcription of common target genes involved in the response to hypoxia. However, stable activation of HIF isoforms with prolonged hypoxia can result in the predominance of HIF-2α activity rather than HIF-1α.^27^ To characterize the contribution of each HIF isoform to these specific pathways in *Vhl*-deficient AMs, we compared the core gene expression signature associated with the response to hypoxia KEGG pathway. The upregulated genes found in *Vhl*-deficient AMs were restored by further deletion of *Hif1a* but not *Hif2a* (Figure S2E). The same was true for DEGs associated to oxidative phosphorylation (OxPhos) and Glycolysis pathways (Figure S2F-G), suggesting that HIF-1α is the main regulator of these genesets over HIF-2α in AMs. We also found that deleting *Hif2a* in *Vhl*-deficient AMs further amplified the transcriptional changes associated to the response to hypoxia, observed in *Vhl* deletion alone (Figure S2F-G). This suggests a negative role of *Hif2a* in the regulation of these pathways. Real-time metabolic profile of AMs in the presence of glucose, pyruvate, and glutamine by Seahorse showed that CD11cΔ*Vhl* and CD11cΔ*Vhl*Δ*Hif2a* AM significantly decreased the basal respiratory rate (BRR) while maintaining extracellular acidification rate (ECAR) compared to controls, reflecting a decreased use of glucose for TCA and mitochondrial oxidative phosphorylation (OxPhos) towards an enhanced aerobic glycolysis and lactate production. In contrast, CD11cΔ*Vhl*Δ*Hif1a* and CD11cΔ*Vhl*Δ*Hif1a*Δ*Hif2a* AM showed levels of BRR and ECAR similar to WT (Figure 1G-H). Overall, our results suggest that the transcriptional program governing AM response to hypoxia, including genes related to OxPhos and glycolysis, is regulated by HIF-1α. Consistent with this, HIF-1α emerges as the key driver of the metabolic switch towards glycolysis in *Vhl*-deficient AMs.

### HIF-1α and HIF-2α differentially regulate AM identity genes in pVHL-deficient mice

We have previously shown that Vhl-deficient AM showed an altered expression of surface markers,^10^ akin to immature cells.^2^ We then asked which HIF isoform dampens mature AM identity in the absence of pVHL. We compared the expression level of classical surface markers that are associated with AM maturation status. While the expression pattern of maturation markers such as CD11c, SiglecF, CD11b or MHC class II (I-A/I-E) was restored to control WT levels in CD11cΔVhlΔHif1aΔHif2a, most of the changes in marker expression found in CD11cΔ*Vhl* mice remained unchanged upon further depletion of HIF-1α. In contrast, these changes were largely reversed upon HIF-2α depletion (Figure 2A-E), suggesting that HIF-2α plays a central role in regulating the impaired AM maturation associated with pVHL loss. We then compared the unique AM gene expression signature that was identified by the Immgen consortium. ^28^ Notably, we found three distinct sets of genes: (i) genes regulated by HIF-2α, with functions in lipid metabolism (Lsr, Vstm2a, Cidec), (ii) genes regulated by HIF-1α, involved in a plethora of functions including regulation of cytoskeleton (Clmn, Sema3e, Krt19, Matn2, Itgax), circadian clock genes (Ear1, Rara) and cell metabolism (Lepr, Naaa, Acaa1b, Aldoc, Fabp1); and (iii) a third set of genes regulated by both HIF-1α and HIF-2α (Figure 2F). Overall, our data indicate that while deleting HIF-2α in Vhl-deficient AMs fully restores the control WT surface marker expression, both HIF isoforms contribute either individually or together shape the subjacent transcriptional identity of mature AMs (Figure 2F).

**Figure 2.**
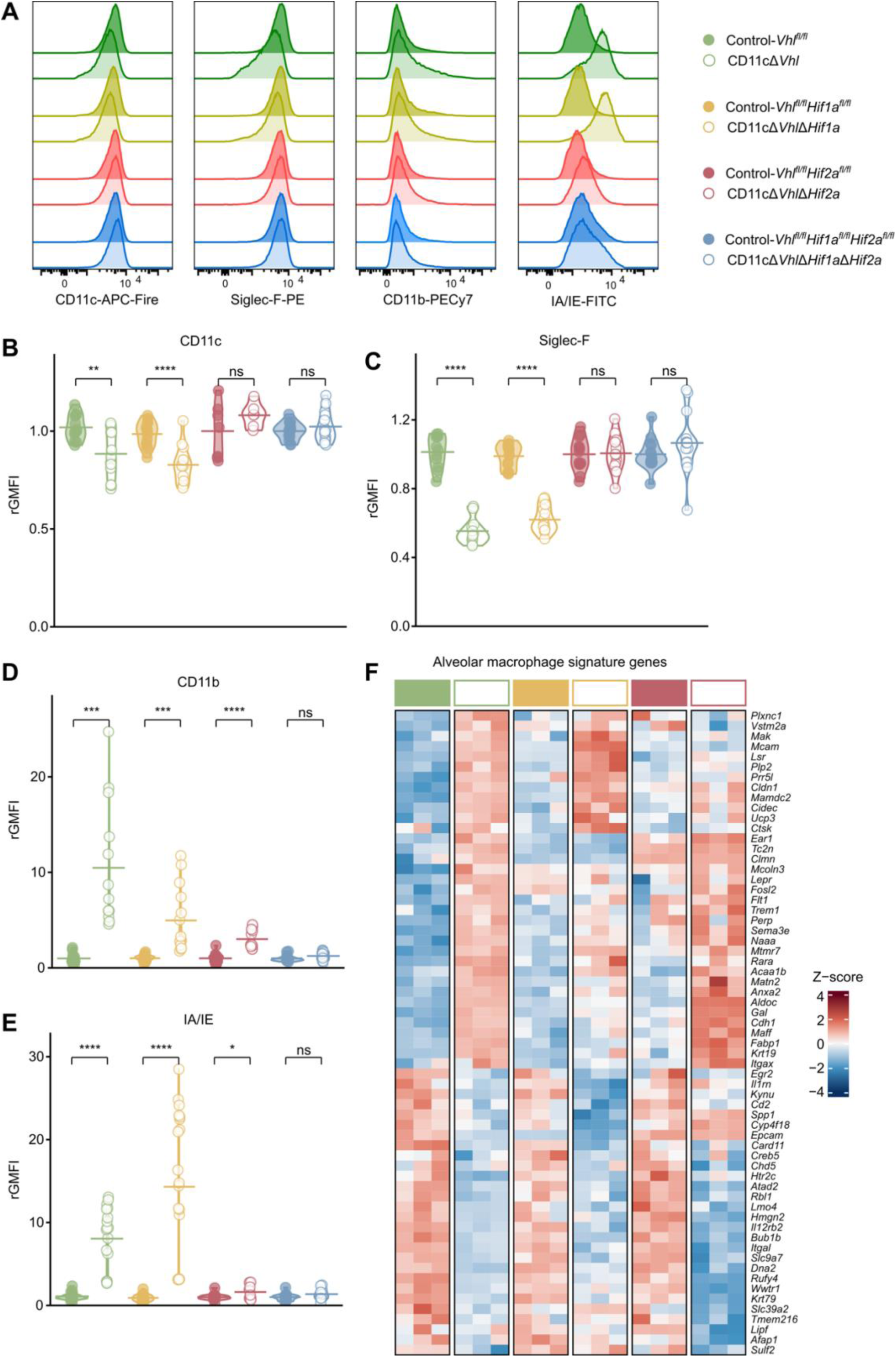
Mature AM transcriptional signature relies on HIF inactivation. (A) Representative flow cytometry histograms for CD11c, Siglec-F, CD11b and IA/IE expression by BAL AMs from the indicated genotypes. (B-E) Quantification of geometric mean of fluoresce intensity (GMFI) relative to the mean value of control samples (rGMFI) of CD11c (B), SiglecF (C), CD11b (D) and IA/IE (E). Pull of 2-3 experiments, n = 4-6 mice per experiment. Student’s *t*-test, ****p<0.0001, ***p<0.001, **p<0.01 and “ns” stands for non-statistically significant. (F) Gene expression heatmap of the lung AM-specific signature gene set described in.^28^

### AM self-renewal capacity is impaired by both HIF-1α and HIF-2α in pVHL-deficient mice

AMs are lung resident cells maintained locally by self-renewal in steady-state.^29^ Enhanced glycolysis has been correlated with reduced AM self-renewal capacity,^5,10,26^ but its association with specific HIF-1 isoforms remains unclear. Thus, we next investigated which HIF isoform was regulating AM proliferation in *Vhl*-deficient AMs. The frequency of Ki67^+^ AMs was significantly and similarly reduced in CD11cΔ*Vhl*Δ*Hif1a* and CD11cΔ*Vhl*Δ*Hif2a* AMs, indicating that both isoforms contribute to the defective proliferation in the absence of pVHL (Figure 3A). In contrast, it was completely restored to control WT in CD11cΔ*Vhl*Δ*Hif1a*Δ*Hif2a* AMs (Figure 3A). In agreement, the differential expression of genes associated to cell cycle in CD11cΔ*Vhl* AM was not restored by the additional single deletion of *Hif1a* or *Hif2a*, being reverted only in CD11cΔ*Vhl*Δ*Hif1a*Δ*Hif2a* AMs (Figure S3A). Furthermore, BrdU uptake in *ex vivo* AM cultures with GM-CSF show a recovery of their full proliferative capacity only in the *Vhl/Hif1a/Hif2a* triple knockout (Figure 3B). Overall, these results demonstrate that HIF-1α and HIF-2α impair AM stemness independently of glycolysis and of environmental extrinsic factors. AMs can differentiate from circulating monocytes coming from bone marrow precursors.^29^ We then asked whether the previously observed compromised AM proliferative capacity constitutes a selective disadvantage at reconstituting an empty niche in a context of competence with WT AM. To address this, we generated bone marrow mixed chimeras: lethally irradiated CD45.1^+^ recipients were transferred intravenously with bone marrow cells from CD45.1^+^CD45.2^+^ WT control and CD45.2^+^ CD11cΔ*Vhl*, CD11cΔ*Vhl*Δ*Hif1a*, CD11cΔ*Vhl*Δ*Hif2a*, CD11cΔ*Vhl*Δ*Hif1a*Δ*Hif2a* or their control Flox littermates in a 1:1 ratio (Figure 3C). After 1.5 months, similar numbers of total AMs were found in all the chimeric mice, ruling out differences in niche accessibility (Figure S3B). Frequencies of blood neutrophils and monocytes from CD45.1^+^CD45.2^+^ and CD45.2^+^ mice were also similar, indicating an equally efficient bone marrow reconstitution (Figure S3C,D). In the bronchoalveolar lavage (BAL), CD45.2^+^ CD11cΔ*Vhl* and CD11cΔ*Vhl*Δ*Hif2a* monocyte-derived AM were significantly less abundant than CD45.1^+^CD45.2^+^ control WT cells (Figure 3D), correlating with a decreased frequency of Ki67^+^ AMs (Figure 3E,F). Unexpectedly, CD11cΔ*Vhl*Δ*Hif1a* monocyte-derived AM also showed a significantly reduced percentage of Ki67^+^ cells but reconstituted the alveolar niche as efficiently as control WT AM, suggesting a survival advantage (Figure 3D-F). These results indicate that while both HIF isoforms regulate AM self-renewal capacity, HIF-1α overexpression may primarily underlie impaired alveolar niche reconstitution upon pVHL absence.

**Figure 3.**
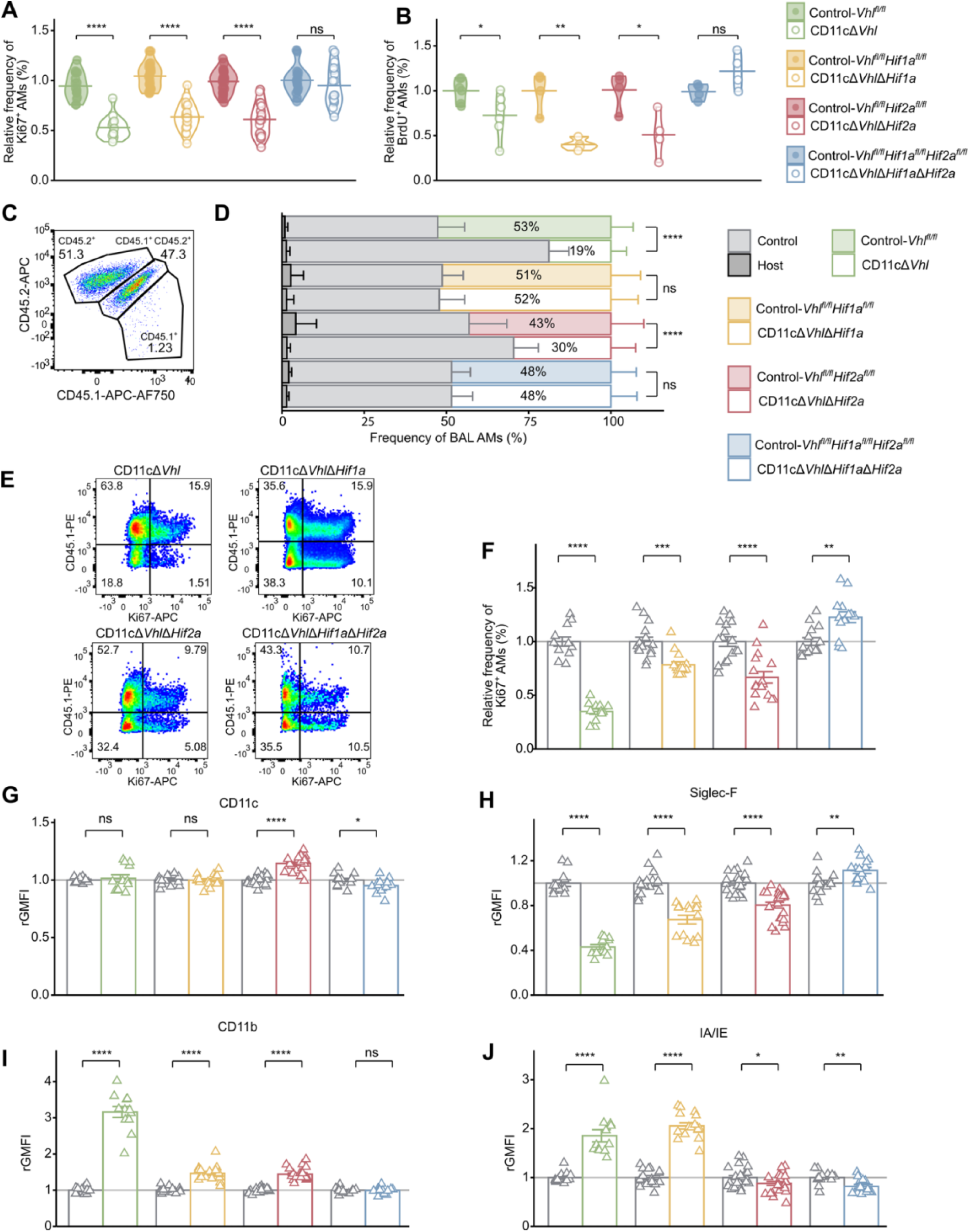
Both HIF isoforms interfere with AM self-renewal. (A) Quantification of the frequency of Ki67+ BAL AMs relativized to the mean value for WT. Pull of 3 experiments, n= 5-8 mice per experiment. Student’s *t*-test, ****p<0.0001, and “ns” stands for non statistically significant. (B) Quantification of the frequency of BrdU+ BAL AMs after 3 days in culture with rGM-CSF. Pool of 2 experiments, each dot represents a pool of n = 15-18 mice per experiment. Student’s *t*-test, **p<0.01, *p<0.05 and “ns” stands for non statistically significant. (C) Representative dot plot showing the percentage of host CD45.1^+^, control transplanted CD45.1^+^CD45.2^+^ and experimental transplanted CD45.2^+^ cells, gated within SiglecF^+^ CD11c^+^ BAL AMs (D) Quantification of the percentage of each AM subpopulation. Bar indicates mean ± SEM, Student’s *t*-test, ****p<0.0001 and “ns” stands for non statistically significant. (E) Representative dot plots showing Ki67 expression by CD45.2^+^ SiglecF^+^CD11c^+^ gated BAL AMs 45 days after BM transfer, in which CD45.1^+^ and CD45.1^-^ correspond to control and experimentally transplanted, respectively. (F) Quantification of the frequency of Ki67+ AMs of each genotype relativized to the mean value for control CD45.1^+^ AMs. Bar indicates mean ± SEM, Student’s *t*-test, ****p<0.0001, **p<0.01. (G-J) Quantification of geometric mean of fluoresce intensity (GMFI) relative to the mean value of control CD45.1^+^ AMs samples (rGMFI) of CD11c (G), Siglec-F (H), CD11b (I) and IA/IE (J). Bar indicates mean ± SEM, Student’s *t*-test, ****p<0.0001, **p<0.01, *p<0.05 and “ns” stands for non statistically significant. (D-J) Pool of experiments, n = 6-8 chimeras per experiment. See also Figure S3.

In this bone marrow-graft model, CD11cΔ*Vhl*, CD11cΔ*Vhl*Δ*Hif1a*, and CD11cΔ*Vhl*Δ*Hif2a* monocyte-derived AMs expressed similar levels of CD11c (Figure 3G) but did not completely restore CD45.1^+^CD45.2^+^ WT control levels for Siglec-F, CD11b, and MHC class II (I-A/I-E) (Figure 3H-J). In contrast, AMs from WT littermates showed frequencies of Ki67^+^ and levels of surface markers similar to CD45.1^+^CD45.2^+^ WT controls (Figure S3E-I). These results confirm that the HIF-dependent defect in the maturation and proliferation of *Vhl*-deficient AM is cell-intrinsic.

### HIF-2α selectively compromises surfactant clearance capacity by VHL-deficient AMs

In lung alveoli, AMs play a major role degrading the surfactant produced by lung type II epithelial cells. Genetic defects in GM-CSF signaling prevent AM maturation and lead to the development of spontaneous PAP, characterized by the accumulation of surfactant.^30^ HIF-1α mediated switch to glycolysis has been linked to the accumulation of lipids and the development of foam cells. ^31–33^ We previously reported that CD11cΔ*Vhl* AMs have a decreased capacity to oxidize lipids *ex vivo*, and consequently, when adoptively transferred to *Cfsr2*^-/-^ mice, which lack AMs and develop PAP spontaneously, they are not able to reduce the surfactant content in the alveoli.^10^ To interrogate on the regulation of lipid metabolism by HIF-1α or HIF-2α *in vivo*, we intratracheally transferred AMs from CD11cΔ*Vhl*Δ*Hif1a*, CD11cΔ*Vhl*Δ*Hif2a*, CD11cΔ*Vhl*Δ*Hif1a*Δ*Hif2a*, or their control counterparts into *Cfsr2*^-/-^ mice. Only adoptively transferred CD11cΔ*Vhl*Δ*Hif2a* AMs and CD11cΔ*Vhl*Δ*Hif1a*Δ*Hif2a* AMs were able to decrease total protein and specific SP-D protein in BAL to levels similar to WT (Figure 4A,B), indicating that HIF-2α stabilization impairs surfactant clearance by lung AMs. Similar numbers of AMs were recovered from BAL for all genotypes, indicating the same efficiency at replenishing the lung alveoli (Figure S4A).

**Figure 4.**
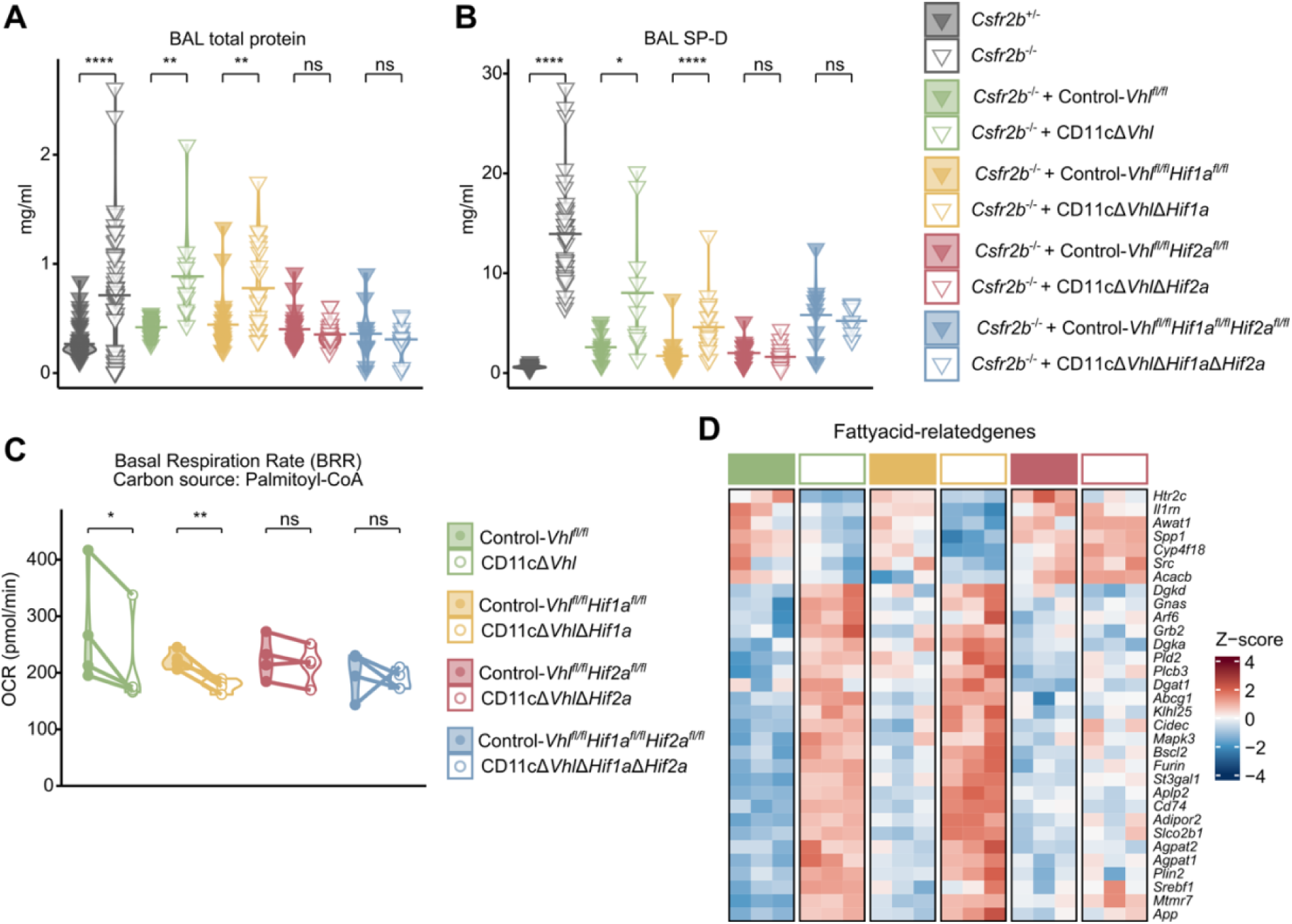
HIF-2 impairs fatty acid oxidation and surfactant clearance by AMs. Quantification of total protein (A) and surfactant protein D (SP-D) (B) in BAL fluid from Csfr2b- /- mice transplanted with BAL AMs from the indicated genotypes. Control mice (Csfr2b^−/−^ and Csfr2b^+/−^) are pooled from each independent experiment. Pool of 3 experiments, n= 3-10 mice per experiment, Student’s t-test, ****p<0.0001, **p<0.01, *p<0.05 and “ns” stands for non statistically significant. (C) Quantification of basal respiratory rate (BRR) of BAL AMs in media containing palmitoyl-CoA and L-carnitine. Pool of 4 experiments, each dot represents a pool of n=6-7 mice. Student’s t-test, **p<0.01, *p<0.05 and “ns” stands for non statistically significant. (D) Heatmap showing differential expression of genes related to lipid metabolic process GO pathway (GO:0019216) in the indicated genotypes. See also figure S4.

AMs rely on fatty-acid oxidation and cholesterol efflux to metabolize lipids and avoid their accumulation respectively.^5^ To assess the regulation of lipid oxidation by HIF isoforms in a controlled *in vitro* setting, we conducted real-time metabolic profile with the fatty acid palmitoyl-CoA as the only carbon source. BRR was significantly reduced in CD11cΔ*Vhl* and CD11cΔ*Vhl*Δ*Hif1a* AMs compared to WT but it was completely restored in CD11cΔ*Vhl*Δ*Hif2a* and CD11cΔ*Vhl*Δ*Hif1a*Δ*Hif2a* AMs (Figure 4C).

We then compared the expression of fatty acid and cholesterol metabolism related genes among genotypes. In agreement with our *in vivo* and *in vitro* results, we found the expression of multiple DEG associated with FA oxidation in *Vhl*-deficient AMs were commonly found in CD11cΔ*Vhl*Δ*Hif1a* but WT expression profile was restored in CD11cΔ*Vhl*Δ*Hif2a* AM (Figure 4D). Globally these results highlight HIF-2α is a major regulator of FA oxidation in lung AMs.

## DISCUSSION

AMs are derived from embryonic progenitors that colonize the lung alveoli and mature after birth. Maturation onset coincides with the exposure of the alveoli to high oxygen concentration upon the first breath. We previously showed that genes related to the adaptation to hypoxia, including HIF-target genes, are downregulated during AM maturation.^10^ The role of HIF-1α is well studied in monocyte-derived macrophages, where it plays a major role in macrophage infiltration and proinflammatory responses.^34–38^ Chronic and acute lung disease due to influenza infection courses with lung edema and cellular infiltrations that can lead to hypoxia. ^39,40^ HIF-1α stabilization has been reported in lung AMs upon lung injury associated to influenza infection, leading to glycolysis induction.^41^ However, the role of the HIF isoforms in AM metabolic adaptation to their niche in homeostasis is largely unknown. In the present study, we have uncovered that HIF-1α and HIF-2α differentially impair specific AM functions. Thus, the inactivation of HIF function is essential for lung AM functional maturation. Our results demonstrate that HIF-1α is the exclusive driver of the glycolytic switch in Vhl-deficient AMs. Notably, both HIF-1α and HIF-2α play a role in the phenotypic maturation and transcriptional identity of AMs, corroborated by the cell-intrinsic nature of the altered surface marker expression found in monocyte-derived AM deficient for Vhl, Vhl/Hif1a and Vhl/Hif2a. HIF-1α stabilization leads to the expression of the antimicrobial enzyme iNOS ^42^ and is protective against lung fibrosis.^26^ This suggests that HIF activation in response to certain exogenous stimuli can be beneficial, whereas under sterile conditions, HIF activity must be tightly regulated.

Increased glycolysis has been linked with a decreased self-renewal potential in AMs.^5,43^ Ki67 expression and BrdU uptake assays indicated that both HIF-1α and HIF-2α stabilization similarly impair AM proliferation in the absence of pVHL. Despite the significant defect in proliferation of *Vhl*-deficient AM, total numbers in the lung alveoli were similar to WT, suggesting that they can fully replenish the alveolar niche in the absence of competition from incoming blood monocytes. Indeed, our experiments with mixed bone marrow chimeras confirmed that that *Vhl*-deficient AM defective proliferation is cell-intrinsic and that *Vhl*- and *Vhl*/*Hif2a*-deficient AMs are significantly outcompeted by control WT AM. In contrast, *Vhl*/*Hif1a*-deficient AMs were similarly able to repopulate the alveoli despite decreased Ki67 expression compared to control WT AM, what could be related with the pro-survival role of HIF-1α ^41^ although further investigation is required.

Notably, we found that compared to *Vhl/Hif1a*-deficient AMs, *Vhl/Hif2a*-deficient AMs had impaired oxidative capacity in the presence of glucose and glutamine as carbon source. In contrast, *Vhl/Hif1a*-deficient AMs showed a decreased oxygen consumption when using FA as carbon source, a phenomenon not observed in Vhl/Hif2a deficient AMs. Investigation on the specific roles of HIF-1α and HIF-2α in the regulation of lipid metabolism pathways is still controversial. HIF-2α has a dominant role leading to hepatic steatosis through fatty acid-oxidation impairment upon depletion of pVHL.^44^ In contrast, HIF-1α was reported to enhance hepatic lipid accumulation in a model of alcoholic fatty liver disease.^32^ Finally, another study shows that deletion both HIF isoforms is sufficient to ameliorate the lipid accumulation induced by hypoxia in HepG2 cell line.^45^ The discrepancy among these results may rely on the different trigger used to activate HIF (pVHL depletion, alcohol and hypoxia, respectively), which likely involves a different dynamic of HIF isoform activation.^27^ In the particular case of macrophages, the source of these cells may determine their metabolic requirements: monocyte derived macrophages (MDM) under hypoxia accumulate lipids and become foam cells, which correlates with the activation of HIF-1α.^31^ HIF-1α can prevent cholesterol efflux contributing to lipid accumulation in a macrophage cell line.^33^ However, MDM strongly rely on glycolysis compared to lung AM at a basal level,^46^ suggesting that the metabolic regulation of MDMs cannot be extrapolated to tissue resident macrophages. In agreement, a recent study has shown that lung AMs are enriched in a fatty acid oxidation module including enzymes involved in lysosomal lipolysis, mitochondrial FA import and oxidation,^47^ Our results unveil HIF-2α as the key regulator of lipid oxidation in lung AMs. *Vhl*/*Hif2a*-deficient AMs completely restore homeostatic levels of surfactant upon transfer to a mouse model of PAP and restore basal OCR levels when using Palmitoyl-CoA as substrate *in vitro*. In agreement, our transcriptional data shows multiple FA-related genes dysregulated in *Vhl*-deficient AMs, which were only restored upon further *Hif2a* inactivation. Overall, this work demonstrates the specific roles of HIF-1α and HIF-2α at fine-tuning lung AM gene expression and functions. Our results highlight the importance of developing therapies targeting each HIF isoform and unveil HIF-2α as a potential target to improve surfactant removal in secondary PAP diseases.

## MATERIALS AND METHODS

### Data

RNA-seq data from Control-*Vhl* and CD11cΔVhl was previously published under GEO: GSE108975 accession number. RNA-seq performed in this study has been deposited at GEO and will be publicly available as of the date of publication (accession number).

### Mice

Mice colonies were bred at Centro Nacional de Investigaciones Cardiovasculares Carlos III (CNIC) under specific pathogen-free conditions. Experiments were conducted with 8- to 10-weeks-old age-matched littermates (regardless of gender). Floxed *Vhl* mice were backcrossed with or without floxed *Hif1a* and floxed *Hif2a*. Additionally, all possible combinations were backcrossed with CD11c-Cre mice (B6.Cg-Tg(*Itgax*-cre)1-1Reiz/J) (kindly provided by Boris Reizis) ^48^ in order to allow the targeting of the alveolar macrophage compartment. *Csf2rb*^-/-^ mice (kindly provided by Martin Guilliams),^49^ and F1 from the mating between C57BL/6J mice expressing the CD45.2 allele (Charles River) and B6/SJL (Ptprca Pepcb/BoyJ) mice expressing the CD45.1 allele were also used. All experimental procedures were approved by the institutional care and use committees and conformed to EU Directive 2010/63EU and Recommendation 2007/526/EC regarding the protection of animals used for experimental and other scientific purposes, enforced in Spanish law under Royal Decree 53/2013.

### AM isolation

Mice were sacrificed with a lethal dose of pentobarbital (Dolethal, Vetoquinol). To isolate AMs for cell culture, ten bronchoalveolar lavages (BAL) were done by inserting a venal catheter (18G BD Pharmingen) into the trachea with 1ml of phosphate-buffered saline (PBS), 5 mM EDTA, and 2 % inactivated fetal bovine serum (FBS, Sigma-Aldrich) (FACS buffer) at 37°C. Samples were kept on ice until further processing. BAL was centrifuged at 300xg for 10’ at 4°C and red blood cell lysis was performed at room temperature (RT) for 3 min with RBC Lysis Buffer (Sigma-Aldrich). Cells were resuspended in medium or FACs buffer depending on the application.

### AM *in vitro* culture

AMs from BAL were cultured in RPMI 1640 (Gibco) supplemented with 10% FBS, 100 U/ml Penicillin and Streptomycin (Lonza), 1mM Sodium Pyruvate (HyClone™), and 2 mM L-glutamine (complete RPMI) at 37°C and 5% CO_2_. For *in vitro* proliferation assay, cells were supplemented with 10 pg/ml rGM-CSF (Peprotech). AMs were pulsed twice with BrdU (BD Pharmingen) at 10μM: 10-12 h and 24h after culture onset. 24h after the second pulse, cells were washed once with PBS to remove remaining FBS and incubated with accutase (StemCell Technologies) for 15-20’ at 37°C. Cells were carefully detached by pipetting. After centrifugation at 600 G for 5’ at 4°C, cells were resuspended and counted, and the same cell number per condition and genotype was stained for surface markers and BrdU.

### Flow cytometry

Single-cell suspensions were stained in 96-U-bottom plates. <u>Surface staining</u> was performed in FACS buffer: PBS, 2.5 mM EDTA and 2.5% heat-inactivated FBS (iFBS). Staining was performed for 15-20’ at 4°C with FcR block anti-mouse CD16/CD32 (2.4G2, Tonbo Biosciences) and the appropriate mix of the following fluorochrome-conjugated antibodies in FACS buffer: CD11c (clone HL3, BD Biosciences), Siglec-F (clone E50-2440, BD Biosciences), CD64 (clone X54-5/7.1, BD Biosciences), CD11b (clone M1/70, BD Biosciences), F4/80 (clone BM8, Life Technologies), MHCII (clone M5/114.15.2 BD Biosciences), Ly6G (clone 1A8, BD Pharmingen), Ly6C (clone AL21, BD Biosciences), CD45 (30-F11, eBiosciences), CD45.1 (A20, eBiosciences), CD45.2 (clone 104, BD Biosciences). Hoechst 33258 (Invitrogen) or Ghost Dye Red 780 (Cell Signaling) were used as a counterstain to exclude dead cells. For <u>intracellular Ki67</u> <u>staining</u>, AMs were stained for surface markers as described above, washed twice with FACs buffer at 600 G for 5’ at 4°C, and mixed with unstained thymocytes, used as a cell carrier. Fixation and permeabilization of cells were performed using the Foxp3/Transcription Factor Staining Buffer Set (#00-5523-00, Invitrogen) and following manufacturer’s instructions. Briefly, fixation solution was prepared by diluting Fixation/Permeabilization concentrate, 1:3 with Fixation/Perm diluent. Permeabilization buffer 10x was diluted 1:10 with sterile water and kept on ice. Cells were fixed for 30’ at 4°C in the dark. Cells were washed with permeabilization buffer 1x and intracellular (i.c.) staining was done right after. Intracellular staining with anti-Ki67-eFluor 660 (SolA15, eBiosciences) or isotype control Rat IgG2aκ (eBR2a, eBiosciences) was performed in permeabilization buffer 1x. All centrifugations after fixation were done at 1000 G for 5’ at 4°C. Cells were resuspended in FACS buffer before acquisition. <u>Intracellular BrdU staining</u> was done following manufacturer instructions. In brief, AMs were stained and mixed with carrier cells as above-mentioned and fixed and permeabilized using the BrdU staining set (BD Pharmingen). Cells were treated with DNAse and stained with anti-BrdU Ab (BD Pharmingen). Samples were acquired in SP6800 Spectral, LSRFortessa, or FACSymphony cell analyzers, and data was analyzed using FlowJo Version 10 software (Tree Star).

### Real-time oxygen consumption rate and glycolytic flux

Real-time oxygen consumption rate (OCR) and extracellular acidification rate (ECAR) were determined with Seahorse XF-96 Extracellular Flux Analyzer (Agilent Technologies). AMs from fresh BAL were pooled from 5-10 mice per genotype. After red blood cell lysis, 2.5x10^5^ AMs were plated in Cell-Tak (Corning) coated wells. The assay was performed in DMEM supplemented with 100 μg/ml penicillin, 100 μg/ml streptomycin, and either 25 mM glucose, 1 mM pyruvate, and 2 mM glutamine or 5 mM L-carnitine and 50 μM palmitoyl-CoA (all Sigma). pH was adjusted to 7.4 with KOH. Three consecutive measurements were performed under basal conditions and after the sequential addition of the following inhibitors: 1 μM Oligomycin, 1 μM Carbonyl cyanide 3-chlorophenylhydrazone (CCCP), and 1 μM Rotenone with 1 μM Antimycin A. Basal respiratory rate (BRR), maximal respiratory rate (MRR), spare respiratory capacity (SRC) and basal ECAR were calculated according to Seahorse Agilent indications.

### RNA sequencing and analysis

CD11c cell positive selection was performed on BAL suspensions from 3-5 mice per genotype. Mouse CD11c MicroBeads UltraPure and MS columns (all Miltenyi Biotec) were used according to manufacturers’ instructions. Briefly, cells were resuspended in 100 μL of FACS buffer plus 20 μL of CD11c MicroBeads, and incubated on ice for 30 mins. Then, cells were washed and purified by magnetic positive selection with MS columns (Miltenyi). A total of 3 pools of 3-5 mice per genotype were included. Total RNA was isolated using Rneasy Micro Kit (Qiagen), and RNA quality was validated with a 2100 Bioanalyzer (Agilent). RNA sequencing was performed in the Genomics Unit of the CNIC on Illumina HiSeq 2500 System. Sequencing reads were pre-processed through a pipeline that used FastQC to assess read quality and Cutadapt to trim sequencing reads, eliminate Illumina adaptor remains, and discard reads that were shorter than 30 base pairs. The resulting reads were mapped against the mouse transcriptome (GRCm38.99) with STAR. Expected expressions were quantified using RSEM, and batch normalization was applied with Combat as implemented in the sva R package. Genes were filtered to have at least 10 CPM, and data were normalized using the trimmed mean of m values (TMM), both with edgeR. DESeq2 rlog-normalized values were used to generate Hexagonal Triwise diagrAMs with Triwise package. Significantly different genes for each pair-wise comparison (p.adj < 0.05, logFC > 1.5) are displayed as red dots in the plots. Genes were ranked according to log-Fold change, and gene set enrichment analysis was performed using FGSEA algorithm and KEGG and GO databases. Redundant GO terms were removed using GOSemSim package. Heatmaps display deregulated genes (Adj. p-value ≤ 0.05) of the specified gene set.

For PCA-based enrichment analysis, log2(CPM) samples were scaled and used as input for principal component analysis (PCA). Genes were ranked according to the loading vectors of PC1. Resulting ranks were used as input for the FGSEA algorithm along with the Kyoto Encyclopedia of Genes and Genomes (KEGG) (www.genome.jp/kegg) and Gene Ontology (GO) (www.geneontology.org/) databases in order to identify the main sources of variability detected by PCA. Only gene sets with a number of genes ≥ 15 and ≤ 500 were considered for PCA-based enrichment analysis.

### Bone Marrow Transfer mixed chimeras

B6/SJL mice expressing the CD45.1 congenic marker were lethally irradiated with 2 doses of 6 Gy that were 3 hours apart and reconstituted intravenously with 5x10^6^ cells of a 1:1 mixture of CD45.1^+^ CD45.2^+^ wild-type BM cells and CD45.2^+^ BM cells coming from the indicated experimental genotypes. Mice were analyzed 45-50 days after reconstitution as previously described. A small sample of peripheral blood was harvested one day before the experimental endpoint to test immune compartment reconstitution.

### Pulmonary macrophage transplantation

5x10^4^ AMs from BAL were transferred intratracheally into Csf2rb^-/-^ mice. Recipient mice were anesthetized with Ketamine (Imalgene, Merial) and Xylazine (Rompun, Bayer). When entirely unconscious, a small incision was made to partially expose the trachea. Then, mice were carefully intubated orotracheally with an IV catheter (22 G; BD Insyte). Cells were inoculated with a pipette in 30μl of PBS. Next, mice were extubated, and the incision sutured. Mice were finally injected with anesthesia reversor (Medeson, Urano) and kept in a warm plaque until complete recovery. 1ml PBS1x-BAL from recipient mice was collected six weeks after transplantation. Cells were centrifuged at 1700 rpm for 5 min at RT and supernatant was used to measure total protein concentration by BCA (Pierce BCA Protein Assay Kit, Thermo Scientific) and surfactant protein D (SP-D) concentration by ELISA (SP-D DUO ELISA kit, R&D System), according to the manufacturer’s instruction. Cell engraftment was compared by flow cytometry of total lung cells.

### Statistical analysis

Statistical comparisons were analyzed using R v.4.2.1. Data are presented as individual values and were analyzed as indicated in the legends or the dedicated methods section. All experiments were repeated at least twice, and pooled data from several experiments are shown as indicated in the legends. All *n* values represent biological replicates (different mice, primary cell preparations, or in vitro experiments). Differences were considered significant when p ≤ 0.05 and are indicated as *ns* when not significant, *p ≤ 0.05, **p ≤ 0.01, ***p ≤ 0.001, ****p ≤ 0.0001.

## ACKNOWLEDGEMENTS

We thank members of the DS laboratory at CNIC, past and present, for feedback and scientific discussions on this project and manuscript. Thanks also to all CNIC facilities and assistants for their support. Work in the DS laboratory is funded by the CNIC; by Ministerio de Ciencia, Innovación y Universidades (MICIU) PID2022-137712OB-I00, CPP2021-008310 and CPP2022-009762 MICIU/AEI/10.13039/501100011033 Agencia Estatal de Investigación, Unión Europea NextGenerationEU/PRTR; by Comunidad de Madrid (P2022/BMD-7333 INMUNOVAR-CM); by Scientific Foundation of the Spanish Association Against Cancer (AECC- PRYGN246642SANC); by Worldwide Cancer Research WWCR-25-0080; by European Union ERC-2023-PoC; by a research agreement with Inmunotek S.L.; and by “la Caixa” Foundation (LCF/PR/HR23/52430012 and LCF/PR/HR22/52420019).

The CNIC is supported by the Instituto de Salud Carlos III (ISCIII), the MICIU and the Pro CNIC Foundation, and is a Severo Ochoa Center of Excellence (CEX2020-001041-S funded by MICIU/AEI /10.13039/501100011033).

