## Supplementary Figures for "HIF-1α and HIF-2α transcription factors differentially regulate lung alveolar macrophage function": 24112025 VHL Priego & Ada╠ün Supp bioRxive.pdf

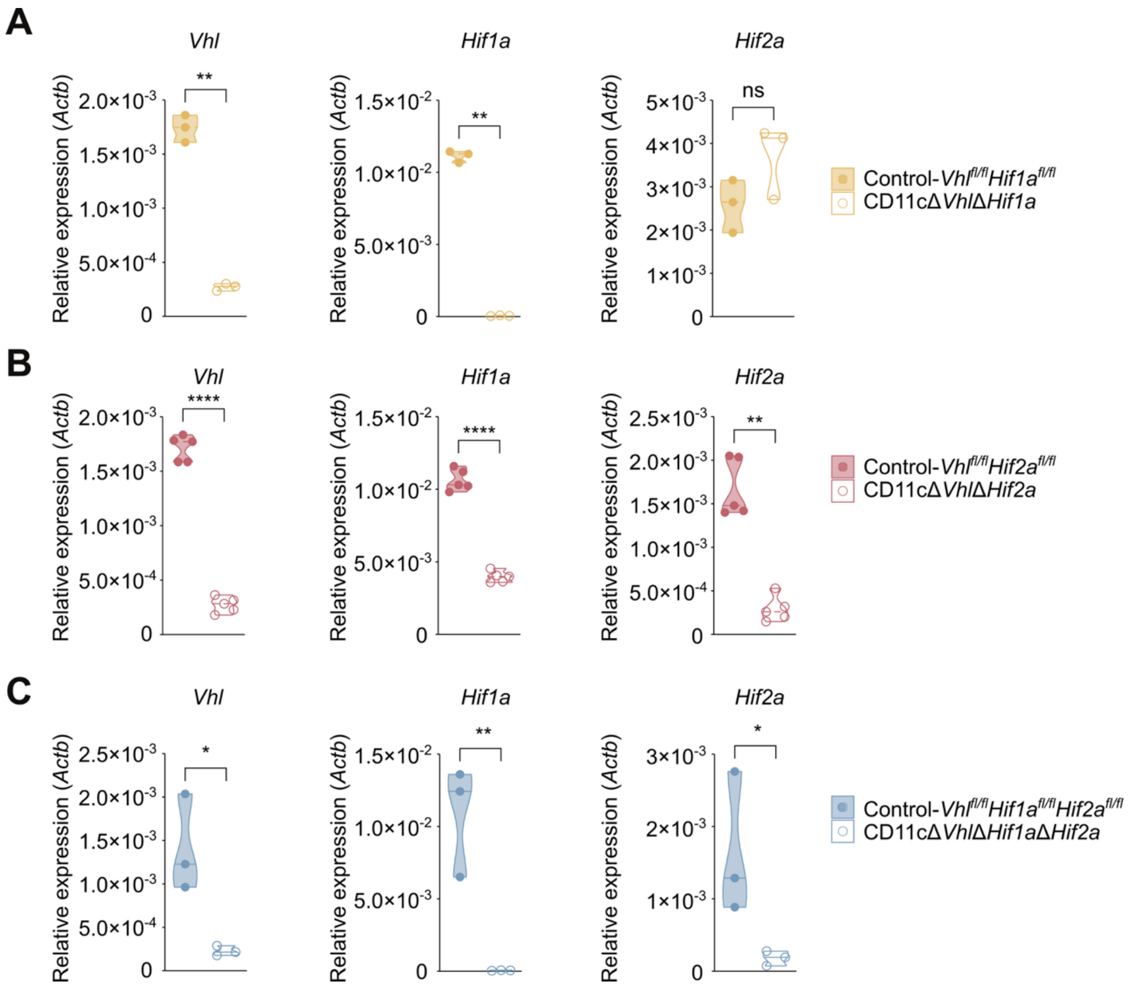


**Figure S1. Validation of *Vhl, Hif1a and Hif2a deletion in* CD11cΔ*Vhl/Hif1a*, CD11cΔ*Vhl/Hif2a, and* CD11cΔ*Vhl/Hif1a/Hif2a* AMs. Related to Figure 1.**

Quantification of the mRNA expression of *Vhl* (left panels), *Hif1a* (middle panels) and *Hif2a* (right panels), normalized to *Actb* mRNA, by BAL AMs from CD11cΔ*Vhl/Hif1a* (A), CD11cΔ*Vhl/Hif2a* (B), and CD11cΔ*Vhl/Hif1a/Hif2a* (C) mice, compared to their control counterparts. Each dot represents a pool of n = 6-7 mice. Paired Student's *t*-test, ****p<0.0001, **p<0.01, *p<0.05 and "ns" stands for non statistically significant.

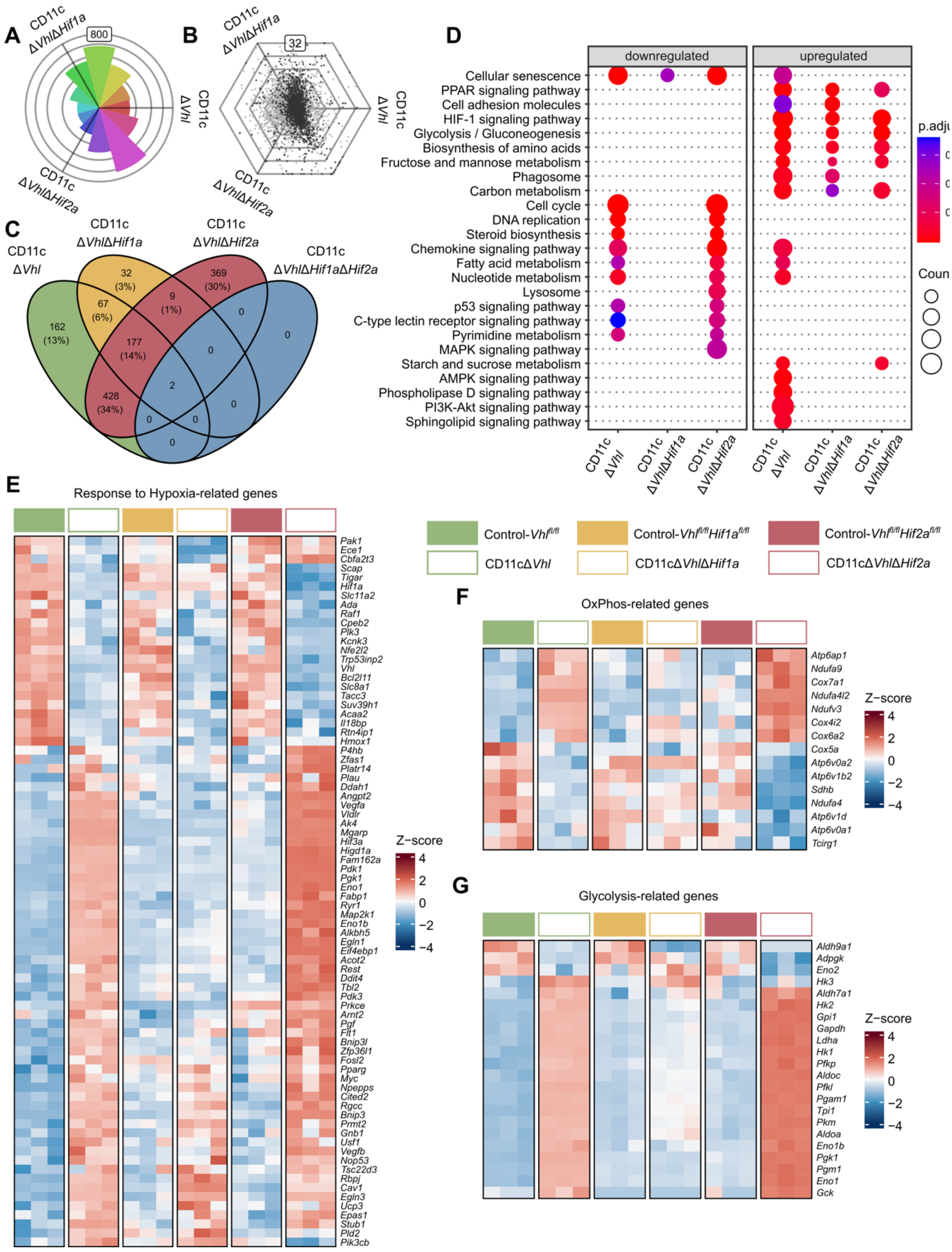


**Figure S2. HIF1α drives glycolysis and adaptation to hypoxia in VHL-deficient alveolar macrophages. Related to Figure 1.**

(A) Rose diagram of all arrayed genes. Frequencies of genes upregulated in one or shared by two genotypes are depicted. (B) Hexagonal diagram showing differentially expressed genes in each genotype. Grid lines represent one $log_2$ fold change. (C) Venn diagram showing shared dysregulated genes between genotypes (FDR-adjusted p-value < 0.05 and

|$\log_2$ fold change| > 1). (D) GO deregulated pathways comparing all genotypes. The size of each dot represents the number of genes, and the color illustrates the pathway enrichment significance according to the adjusted p-value. (E-G) Heatmaps representing Z-scores of differentially expressed genes associated to the response to hypoxia (GO:0001666, E), OxPhos (mmu00190, F) and Glycolysis (mmu00010, G) pathways.

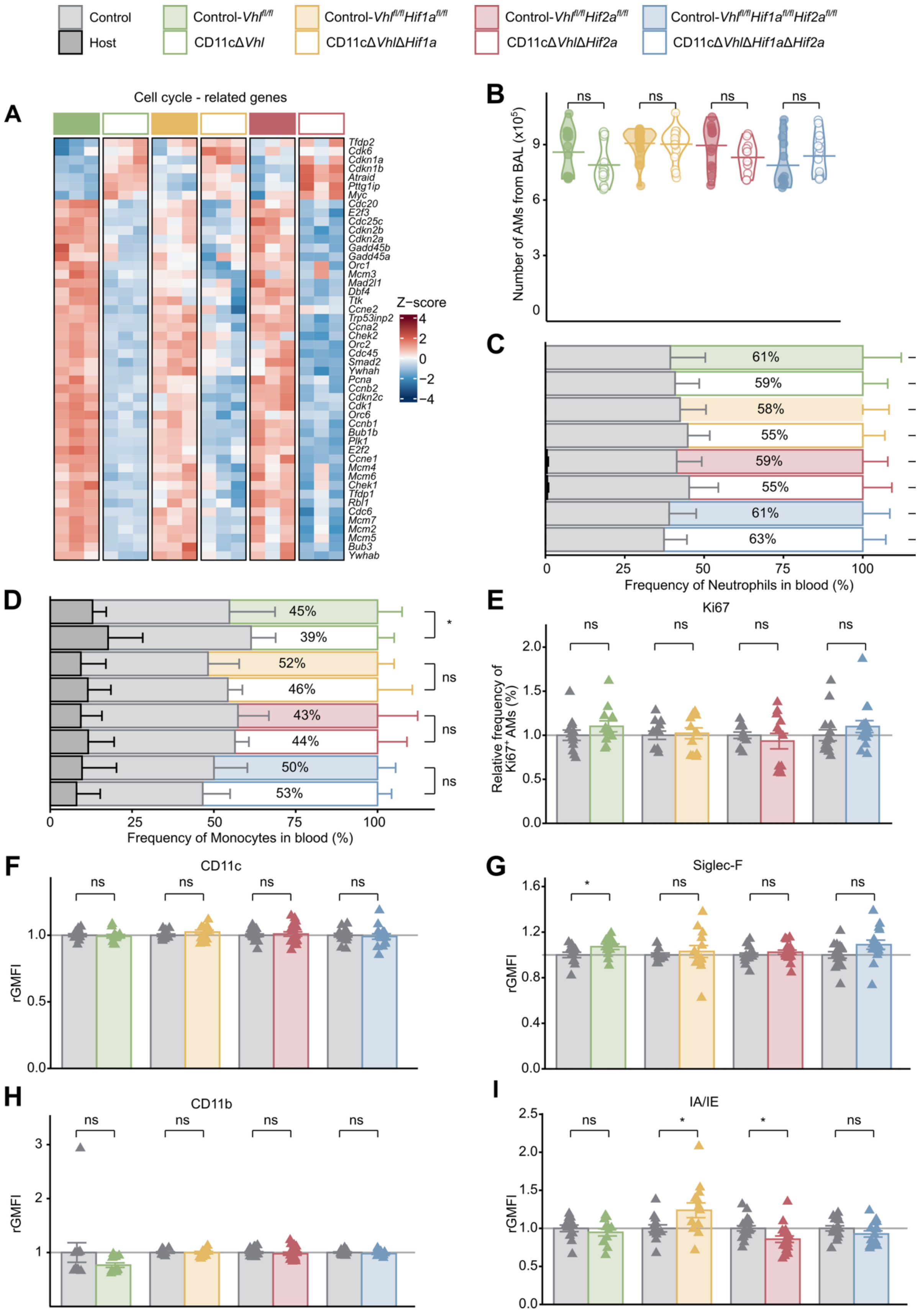


**Figure S3. Reconstitution of bone marrow chimeras. Related to Figure 3.**

(A) Heatmap depicting differentially expressed genes included in the Cell cycle Kegg pathway (mmu04110). (B) Quantification of the number of CD45.2$^{+}$ AMs in BAL from mouse chimeras. Student's *t*-test, "ns" stands for non statistically significant. (C-D) Quantification of the frequency of blood neutrophils gated as $CD11b^{+}Ly6G^{+}H58^{-}$ (C) and blood monocytes gated as $CD11b^{+}Ly6C^{+}H58^{-}$ (D) from host $CD45.1^{+}$, control $CD45.1^{+}CD45.2^{+}$ and $CD45.2^{+}$ in BM chimeras 40 days after transfer. Student's *t*-test, $^{*}p<0.05$ and "ns" stands for non statistically significant. (E) Quantification of the frequencies of $Ki67^{+}$ AMs in control $CD45.1^{+}CD45.2^{+}$ and $CD45.2^{+}$ AMs with the indicated genotype. Student's *t*-test, "ns" stands for non statistically significant. Quantification of geometric mean of fluoresce intensity (GMFI) relative to the mean value of control $CD45.1^{+}$ AMs samples (rGMFI) of CD11c (F), Siglec-F (G), CD11b (H) and IA/IE (I) in $CD45.2^{+}$ AMs with the indicated genotype. Student's *t*-test, $^{*}p<0.05$ and "ns" stands for non statistically significant. (B-I) Pool of 3 experiments, n= 3-10 mice per experiment.

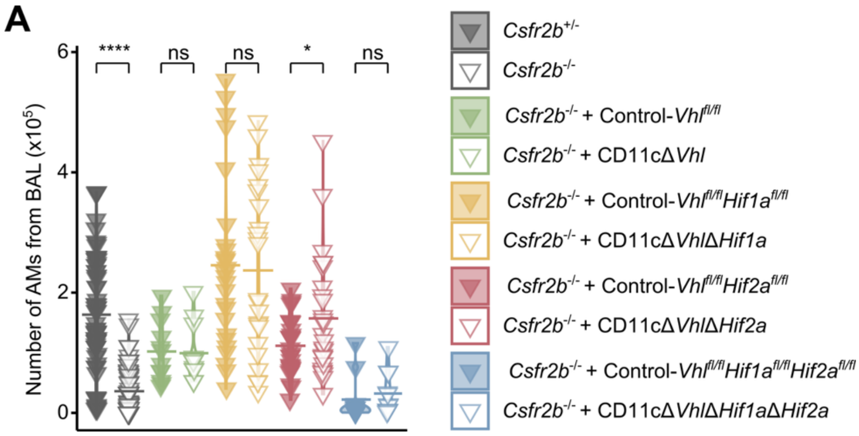


**Figure S4. Lung niche reconstitution by AMs. Related to Figure 4.**

(A) Quantification of the total number of AMs from the lung of Csfr2b$^{-/-}$ mice 45 days after pulmonary macrophage transplantation with BAL AMs from the indicated genotypes. Pool of 3 experiments, n = 3-10 mice per experiment. Student's *t*-test, ****p<0.0001, *p<0.05 and "ns" stands for non statistically significant.
